# A comparative ‘omics approach to candidate pathogenicity factor discovery in the brain-eating amoeba *Naegleria fowleri*

**DOI:** 10.1101/2020.01.16.908186

**Authors:** Emily K. Herman, Alex Greninger, Mark van der Giezen, Michael L. Ginger, Inmaculada Ramirez-Macias, Haylea C. Miller, Matthew J. Morgan, Anastasios D. Tsaousis, Katrina Velle, Romana Vargová, Sebastian Rodrigo Najle, Georgina MacIntyre, Norbert Muller, Mattias Wittwer, Denise C. Zysset-Burri, Marek Elias, Claudio Slamovits, Matthew Weirauch, Lillian Fritz-Laylin, Francine Marciano-Cabral, Geoffrey J. Puzon, Tom Walsh, Charles Chiu, Joel B. Dacks

**Affiliations:** Division of Infectious Disease, Department of Medicine, Faculty of Medicine and Dentistry, University of Alberta; Department of Agricultural, Food and Nutritional Science, University of Alberta; Laboratory Medicine and Medicine / Infectious Diseases, UCSF-Abbott Viral Diagnostics and Discovery Center, UCSF Clinical Microbiology Laboratory UCSF School of Medicine; Department of Laboratory Medicine, University of Washington Medical Center; Centre for Organelle Research, University of Stavanger; School of Applied Sciences, Department of Biological and Geographical Sciences, University of Huddersfield; Department of Cardiology, Hospital Clinico Universitario Virgen de la Arrixaca. Instituto Murciano de Investigación Biosanitaria. Centro de Investigación Biomedica en Red-Enfermedades Cardiovasculares (CIBERCV); CSIRO Land and Water; CSIRO Land and Water, Black Mountain Laboratories; Laboratory of Molecular and Evolutionary Parasitology, RAPID group, School of Biosciences, University of Kent, Canterbury, UK; Department of Biology, University of Massachusetts; Department of Biology and Ecology, Faculty of Science, University of Ostrava; Institut de Biologia Evolutiva (UPF-CSIC); Department of Medicine, Faculty of Medicine and Dentistry, University of Alberta; Institute of Parasitology, Vetsuisse Faculty Bern, University of Bern; Spiez Laboratory, Federal Office for Civil Protection, Austrasse, Spiez, Switzerland; Department of Ophthalmology, Inselspital, Bern University Hospital, University of Bern; Department of Biochemistry and Molecular Biology, Centre for Comparative Genomics and Evolutionary Bioinformatics, Dalhousie University; Center for Autoimmune Genomics and Etiology and Divisions of Biomedical Informatics and Developmental Biology, Cincinnati Children’s Hospital Medical Center, Cincinnati, Ohio, USA, Department of Pediatrics, University of Cincinnati College of Medicine; Department of Microbiology and Immunology, Virginia Commonwealth University School of Medicine; Department of Life Sciences, The Natural History Museum; Institute of Parasitology, Biology Centre, Czech Academy of Sciences

**Keywords:** Illumina, RNASeq, genome sequence, protease, cytoskeleton, metabolism, lysosomal, inter-strain diversity, neuropathogenic

## Abstract

Of the 40 described *Naegleria* species, only *N. fowleri* can establish infection in humans, killing almost invariably within two weeks. In the brain, the amoeba performs piece-meal ingestion, or trogocytosis, of brain material causing massive inflammation. Conversely, its close relative *Naegleria gruberi*, which is used as a laboratory model organism, is non-pathogenic. The exact pathogenicity factors distinguishing *N. fowleri* from its harmless relatives are unclear. We have here taken an -omics approach to understanding *N. fowleri* biology and infection at the system level. We provide the first analysis of genomic diversity between strains, finding little conservation in synteny but high conservation in protein complement. We also demonstrate that the *N. fowleri* genome encodes a similarly complete cellular repertoire to that found in *N. gruberi*. Our comparative genomic analysis, together with a transcriptomic analysis of low versus high pathogenicity *N. fowleri* cultured in a mouse infection model, allowed us to construct a model of cellular systems involved in pathogenicity and furthermore provides ~500 novel candidate pathogenicity factors in this currently rare but highly fatal pathogen.

## Introduction

*Naegleria fowleri* is an opportunistic pathogen of humans and animals, causing primary amoebic meningoencephalitis (PAM), and killing up to 97% of those infected, usually within two weeks (Carter, 1970). It is found in warm freshwaters around the world and drinking water distribution systems (DWDS) (Morgan et al., 2016; Puzon et al., 2009; Puzon et al., 2017) with *N. fowleri*-colonized DWDSs linked to deaths in Pakistan (Kazi and Riaz, 2013; Mahmood, 2015; Naqvi et al., 2016), Australia (Dorsch, 1982), and the USA (Cope et al., 2015; Yoder et al., 2012). Infection occurs when contaminated water enters the nose. *N. fowleri* passes through the cribriform plate to the olfactory bulb in the brain and phagocytoses brain material, causing physical damage and massive inflammation leading to death. Although successful treatment with miltefosine and other antimicrobials is becoming more common (Cope et al., 2016; Linam et al., 2015), this relies on appropriate early diagnosis, which is challenging due to the relatively low incidence of amoebic versus viral or bacterial meningitis.

The reported incidence of infection is relatively low (Trabelsi et al., 2012); however, infection is likely more prevalent, particularly in developing countries with warm climates and inconsistent medical reporting. *N. fowleri* has been recently proposed as an emerging pathogen based on increased case reports in the past decade (MacIver et.al. 2019). With cases from temperate locations reported in recent years (Baral and Vaidya, 2018; Kemble et al., 2012), the threat of *N. fowleri* may be exacerbated by range expansion due to climate change (MacIver et.al. 2019), which has been associated with a rise in freshwater temperatures, an increase in aquatic recreational activities (Siddiqui and Khan, 2014), and extreme weather events. It is likely that we are still not fully aware of the global scale of *N. fowleri* infection (Maciver et al., 2019), or how infection may increase as climate change accelerates.

Of the ~40 species of *Naegleria*, found commonly in soils and fresh waters worldwide, *N. fowleri* is the only species that typically infects humans, suggesting that pathogenicity is a gained function. Several labs have identified potential *N. fowleri* pathogenicity factors including proteases, lipases, and pore-forming proteins. The hypothesized mechanism of *N. fowleri* pathogenicity – tissue degradation for both motility during infection and phagocytosis – is compatible with these factors. However, none appear to be unique to *N. fowleri*, and there are likely proteins factors yet to be identified that are responsible for pathogenesis. In 2010 the genome sequence of the non-pathogenic *Naegleria gruberi* was reported and in 2014, the genome of *N. fowleri* ATCC 30863 was sequenced and used to guide comparative proteomic work on highly and weakly pathogenic *N. fowleri* (Zysset-Burri et al., 2014). The genome of a thermotolerant but non-pathogenic species, *Naegleria lovaniensis*, was published in 2018 (Liechti et al. 2018). Despite such work, a genome-wide perspective on *N. fowleri* diversity and pathogenesis is lacking.

Here we report the genome sequences of two *N. fowleri* strains; 986, an environmental isolate from an operational DWDS in Western Australia, and CDC:V212, a strain isolated from a patient. We also report a transcriptomic analysis of induced pathogenicity in the *N. fowleri* strain LEE to identify the genes differentially expressed as a consequence of infection.

## Results

The genomes of *N. fowleri* strains V212 and 986 were sequenced to average coverage of 251 × and 250 × respectively and the transcriptome of axenically cultured *N. fowleri* V212 was also sequenced to support gene prediction. Analysis of genome architecture shows limited synteny between *N. fowleri* strains but conservation of other genome statistics, as compared to the *N. gruberi* genome (Figure 1, Table 1). Similarly, while GC content and other coding content statistics were similar between *N. fowleri* strains, they were remarkably different from those values for *N. gruberi* (Table 1) (Fritz-Laylin et al., 2010a). The differences in genome statistics are not unreasonable given that genetic diversity within the *Naegleria* clade has been equated to that of tetrapods (Baverstock et al., 1989). Importantly, there is transcriptomic evidence for 82% of genes in *N. fowleri* V212, suggesting that they are expressed when grown in culture and 100% of the genes with transcriptome evidence were contained within the predicted genomic set. Out of 303 near-universal single-copy eukaryotic orthologues, 277 were found in the set of *N. fowleri* V212 predicted proteins, giving a BUSCO score of 91.4%, signifying that the genome and predicted proteome are highly complete.

**Table 1.** Genome statistics for *N. fowleri* strains V212, 986, and ATCC 30863, and *N. gruberi* strain NEG-M.

| *Species* | Total genome size | GC content | Number of genes | Average gene length | Exons/gene | Average exon length | % Coding | Average intron length |
| --- | --- | --- | --- | --- | --- | --- | --- | --- |
| *N. fowleri* V212 | 27.7 Mbp | 36% | 12,677 | 1,785 bp | 2 | 777 bp | 71.35% | 126 bp |
| *N. fowleri* 986 | 27.5 Mbp | 36% | 11,599 | 1,955 bp | 2 | 849 bp | 73.01% | 138 bp |
| *N. fowleri* ATCC 30863 | 29.62 Mbp | 35% | 11,499 | 1,984 bp | 2 | 825 bp | 70.79% | 144 bp |
| *N. gruberi* NEG-M | 41.0 Mbp | 33% | 15,708 | 1,677 bp | 1.7 | 894 bp | 57.8% | 203 bp |

**Figure 1.**
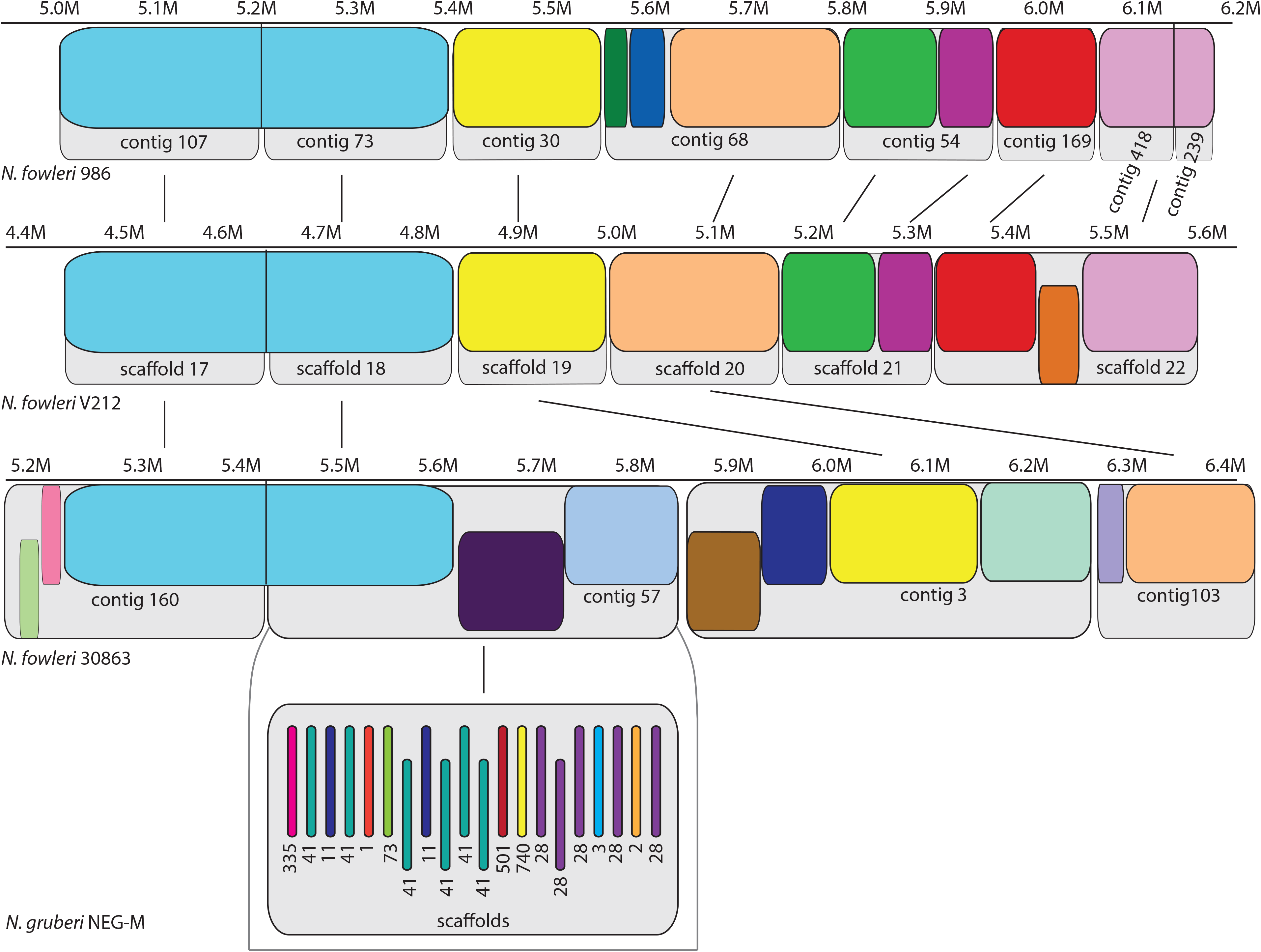
An example of genomic synteny conservation among the three *N. fowleri* isolates. Because of low conserved synteny between *N. fowleri* and *N. gruberi*, the organization of homologous sequence in *N. gruberi* is shown for only one *N. fowleri* contig. Sequence lengths are shown above each genome in millions of bases (M). As the contig and scaffold alignments depicted are a subset representative of the complete alignment of the assemblies, the scale bars above each assembly are relative to the complete alignment. They are included to show the relative sizes of the regions of homologous sequence. Numbering of scaffolds is arbitrary. Blocks of shared sequence are colour-coded, while contig or scaffold divisions are shown as grey boxes. Blocks that are syntenic between strains within the visual space are indicated by lines, while blocks without lines also have homologous sequence elsewhere in the two other strains, but cannot be shown due to the large intervening distance. Offset blocks indicate reverse complemented sequence relative to *N. fowleri* 986.

### *The* Naegleria fowleri *genome encodes a complete and canonical cellular complement*

In 2010, *N. gruberi* was hailed as possessing extensive eukaryotic cytoskeletal, membrane-trafficking, signaling, and metabolic machinery, suggesting sophisticated cell biology for a free-living protist that diverged from other eukaryotic lineages over one billion years ago. Careful manual curation and genomic analysis of meiotic machinery (Supplementary Figure 1, Supplementary Table 1) transcription factors (Supplementary Table 2, Supplementary Material 1), sterols (Supplementary Table 3, Supplementary Material 2, Supplementary Figures 2 and 3), mitochondrial proteins (Supplementary Table 4), cytoskeletal proteins (Supplementary Table 5, Supplementary Figure 4, Supplementary Material 3), membrane trafficking components (Supplementary Table 6), and small GTPases (Supplementary Material 4, Supplementary Figures 5–7, Supplementary Table 7) demonstrates that like *N. gruberi*, *N. fowleri* possesses a remarkably complete repertoire of cellular machinery.

### *A comparative approach to identifying the cellular basis of* N. fowleri *pathogenicity*

As pathogenesis is likely a gain-of-function in *N. fowleri*, we specifically looked for differences with the non-pathogenic *N. gruberi*. Using OrthoMCL, the proteins from the three *N. fowleri* strains and *N. gruberi* were clustered into 11,399 orthogroups, of which 7,656 (67%) appear to be shared by all four *Naegleria* species, and 10,451 (92%) are shared by all three *N. fowleri* strains (Figure 2). There are 2,795 groups not identified in *N. gruberi* that are shared by all three *N. fowleri* strains. Manual verification of orthology reduced the number of orthogroups with genes in *N. fowleri* but not *N. gruberi* to 458 proteins (Supplementary Table 8.) Of these, 80% are unique to *N. fowleri,* with no clearly homologous sequence in other organisms based on NCBI BLAST, and only 52 (11%) identified either could be functionally annotated based on NR BLAST results or contain a characterized domain. 404 of the 458 genes have substantial transcriptomic evidence (FPKM >5), suggesting that most of the gene models are accurate and these genes are expressed.

**Figure 2.**
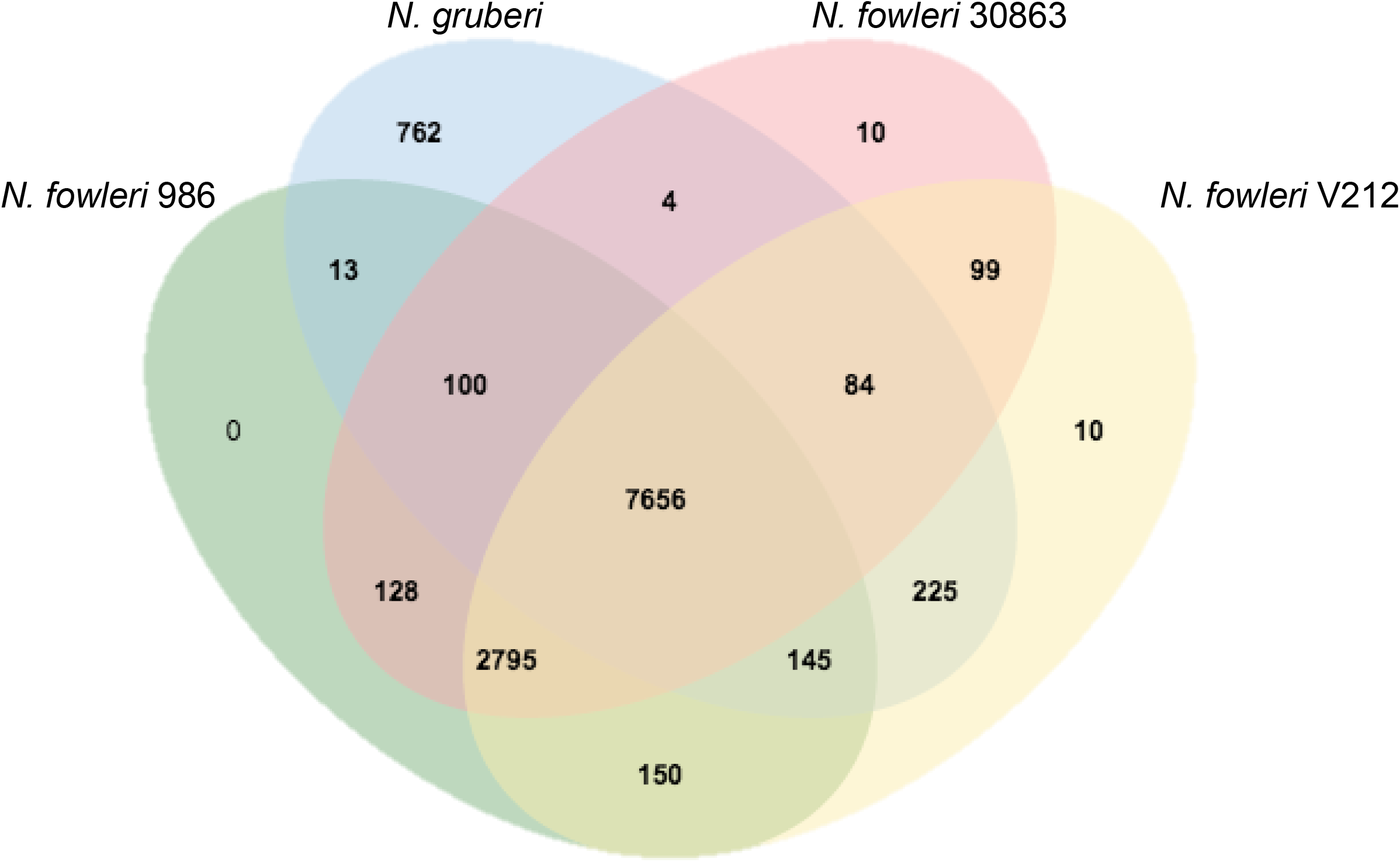
Result of OrthoMCL analysis showing the number of orthogroups shared between the three *N. fowleri* strains and *N. gruberi.* The number of in-paralogue groups within each species is also shown.

In order to prioritize the candidate pathogenesis factors, we took a transcriptomics approach taking advantage of previously established experimentally-induced pathogenicity in the *N. fowleri* LEE strain (Whiteman and Marciano-Cabral, 1987). In this system, not only does mouse passaged *N. fowleri* have a lower LD_50_ in guinea pigs by two orders of magnitude, it is also resistant to complement-mediated killing, while cultured *N. fowleri* LEE and *N. gruberi* are not (Whiteman and Marciano-Cabral, 1987).

In our analysis, there are 315 differentially expressed genes in mouse-passaged *N. fowleri* LEE (LEE-MP) versus *N. fowleri* LEE grown only in culture (LEE-Ax) (Supplementary Table 9). Of these, 206 are up-regulated in high pathogenicity *N. fowleri*, while 109 are down-regulated (Supplementary Figure 8), and in terms of function, span multiple cellular systems with potential links to pathogenesis.

Overall analysis of down-regulated genes was less informative than that of up-regulated genes. Systems represented in the down-regulated gene dataset include signal transduction, flagellar motility, genes found to be involved in anaerobiasis in bacteria and transcription/translation (Supplementary Table 9). Nonetheless, approximately 70% of the down-regulated genes not in these categories are genes of unknown function, and many are specific to *N. fowleri* or *Naegleria* spp.

Building on the manual curation of the encoded cellular machinery and using the comparative genomic and transcriptomic data, we confirmed and extended our knowledge of previously identified categories of pathogenicity factors, and identified several novel major aspects of *N. fowleri* biology with implications for pathogenesis and that might represent avenues for investigation.

### Proteases are leading candidates for pathogenicity factors

*N. fowleri* is known to secrete proteases to traverse the extracellular matrix during infection, and pore-forming proteins to kill and digest host cells (Aldape et al., 1994; Herbst et al., 2002; Hu et al., 1991; Serrano-Luna et al., 2007; Toney and Marciano-Cabral, 1992). Perhaps the most prominent example is a prosaposin pore-forming protein (termed Naegleriapore A), which is heavily glycosylated and protease-resistant (Herbst et al., 2002). Consistent with this, the precursor protein for Naegleriapore A and B was found to be in our up-regulated DE gene set. Similarly, the cathepsin protease (Cathepsin A, or Nf314) is up-regulated in highly pathogenic *N. fowleri*, consistent with it previously being identified as a pathogenicity factor (Hu et al., 1992).

Comparative genomics found broadly similar complements of proteases between the *N. fowleri* strains and *N. gruberi* (Supplementary Figure 9, Supplementary Table 10), with one exception. The serine protease S81 was found in all three *N. fowleri* genomes but not in *N. gruberi*. The only homologue of this protein is found in *Hirudo medicinalis*, the European medicinal leech. This protein has a destabilase and peptidoglycan-binding domain; it is found in the saliva of the leech and it may have lysozyme activity and be capable of dissolving fibrin (Zavalova et al., 1996; Zavalova et al., 2000). The gene was highly (but not differentially) expressed under both axenic and mouse-passaged conditions with FPKM values ranging from 500-800.

Twenty-eight proteases are up-regulated in highly pathogenic *N. fowleri*, making up more than 10% of all up-regulated genes (Supplementary Table 9). Of the protease families with *N. fowleri* homologues up-regulated following mouse passage, half are either localized to lysosomes or are secreted, while the others have proteolytic activities in other organelles or in the cytoplasm. The most substantially represented types of lysosomal/secreted protease in the up-regulated genes are the cathepsin proteases; specifically, the C01 subfamily, with 10 out of 21 genes up-regulated in highly pathogenic *N. fowleri*. The C01 subfamily includes cathepsins B, C, L, Z, and F. Each of these subfamilies has multiple members, up to 10 in the case of cathepsin B, and members of the B, Z, and F subfamilies are up-regulated.

Despite the large number of C01 subfamily cathepsin proteases in *N. fowleri* (20-21 members), *N. gruberi* encodes even more (35). Many of the *N. fowleri* and *N. gruberi* cathepsins have 1:1 orthology (Supplementary Figure 10) with at least three expansions that have occurred in the cathepsin B clade in *N. gruberi*. These expansions account for most of the difference in paralogue number between the two species, making up 12 of ~16 *N. gruberi*-specific C01 homologues. Notably, of the *N. fowleri* cathepsin genes that are up-regulated in highly pathogenic *N. fowleri* at least two lack orthologues in *N. gruberi*, raising the possibility of their specific involvement in pathogenesis. Protease secretion is underpinned by the membrane trafficking system (the canonical secretion pathway) and autophagy-based unconventional secretion (for proteins lacking N-terminal signal peptides). In both cases, there are few differences in gene presence, absence, and paralogue number in the different *Naegleria* genomes (Supplementary Tables 6 and 11). Strikingly, however, 42% of the up-regulated genes are involved in lysosomal processes. In addition to the 22 proteases above, a lysosomal rRNA degradation gene is up-regulated, as well as three subunits of the vacuolar ATPase proton pump (116 kDa, 21 kDa, and 16 kDa) responsible for acidification of both lysosomes and secretory vesicles. Endo-lysosomal trafficking genes Rab GTPase Rab32 (one of three paralogues) and the retromer component Vps35 were up-regulated.

### Proteins driving actin cytoskeletal rearrangements may contribute to pathogenesis

While actin is known to drive many cellular processes in eukaryotes, Nf-Actin has been considered a pathogenicity factor in *N. fowleri* due to its role in trogocytosis via food cup formation (Sohn et al., 2010). Furthermore, actin binding proteins and upstream regulators of actin polymerization were reported to correlate with virulence (Zysset-Burri et al., 2014). While the complements of actin-associated machinery was similar between the Naegleria species, we notably identified a PTEN domain on one of the formin homologues in *N. fowleri,* that was not identified in *N. gruberi* (Supplementary Material 3). Human also do not encode formins of the PTEN family.

Although *Naegleria* actin protein levels do not always correlate with transcript levels (Fritz-Laylin et al., 2010b; Sussman et al., 1984), a single subunit of the Arp2/3 complex (Arp3) and the WASH complex member strumpellin were both up-regulated in the mouse-passaged amoebae (Supplementary Table 5, Supplementary Material 3). Similarly, we identified an up-regulated RhoGAP22 gene and the serine/threonine protein kinase PAK3, which are involved in Rac1-induced cell migration in other species as well as an up-regulated member of the gelsolin superfamily in the mouse-passaged *N. fowleri*, which may contribute to actin nucleation, capping, or depolymerization (Nag et al., 2013). Although the shift from the environment in the mouse brain to tissue culture conditions prior to sequencing may have resulted in an up-regulation in macropinocytosis, which can alter cell motility and the transcription of cytoskeletal genes in *Dictyostelium discoideum* (Kayman and Clarke, 1983; Sillo et al., 2008) potential importance of cytoskeletal dynamics in promoting virulence.

### *Neither LGT nor cell stress appear to be major drivers of* N. fowleri *pathogenesis*

One obvious potential source of pathogenicity factors is lateral gene transfer of bacterial genes into *N. fowleri* to the exclusion of *N. gruberi* and other eukaryotes. However, of the 458 genes exclusive to *N. fowleri* most are of unknown function (Supplementary Table 8) and of these, only 26 have Bacteria, Archaea or virus as the largest taxonomic group containing the top five BLAST hits. Furthermore, only one of the genes up-regulated in highly pathogenic *N. fowleri* lacks a homologue in *N. gruberi*, and it has a potential homologue in *D. discoideum* and members of the Burkholderiales clade of bacteria (NfowleriV212_g4665, Supplementary Table 9).

Another potential reason for *N. fowleri*’s ability to infect humans and animals is the ability to survive the stresses of infection. However, we observed no obvious differences between the *N. fowleri* and *N. gruberi* complements of the ER-associated degradation machinery and unfolded protein response machinery that would suggest a differential ability to cope with cell stress (Supplementary Table 12). Furthermore, our transcriptomics analysis showed a general down-regulation of cell stress systems, as well as DNA damage repair (Supplementary Table 9). This does not suggest that these systems are significantly involved in pathogenesis and our transcriptomics experiment reflects an organism that is not under duress.

### Adhesion factors as candidate pathogenicity factors

Since infection requires the ability to attach to cells of the nasal epithelium, differences between *N. fowleri* and *N. gruberi* in cell-cell adhesion factors may be relevant to pathogenesis. While we were unable to find evidence of a previously reported integrin-like protein (Jamerson et al., 2012) in any of the genomic data (Supplementary Table 13), we did find that another previously identified attachment protein, Nfa1, was highly expressed in both mouse passaged and axenically cultured *N. fowleri* LEE (>1000 FPKM), providing further evidence for its previously reported role as a general pathogenicity factor (Kang et al., 2005).

*N. fowleri* and *N. gruberi* encode relatively few putative adhesion G protein-coupled receptors (AGPCRs), with 10 or fewer in each organism (Supplementary Table 14). These were identified by searching for proteins with the appropriate domain organization: sequences that have both an extracellular domain (assuming correctly predicted topology in the membrane) and seven transmembrane regions. While their specific functions remain unknown, this work provides a list of proteins that can be later characterized by *in vivo* work.

Of the proteins involved in adhesion in *D. discoideum* (TM9/Phg1, SadA, SibA, and SibC), only orthologues of the TM9 protein could be reliably identified (Supplementary Table 15), which was not found to be differentially regulated in our transcriptomic dataset. It is possible that the *Naegleria* homologue may be involved in cell adhesion, but with other downstream effectors. Given the roles for these proteins in both signalling and adhesion in other systems, cell biological work in *Naegleria* is required to confirm their role.

### The metabolic contribution to pathogenicity

Strikingly, 19% of up-regulated genes are involved in metabolism (Supplementary Table 9). Both catabolic and anabolic processes are represented; some up-regulated genes include phospholipase B-like genes necessary for beta-oxidation, and genes involved in phosphatidate/phosphatidylethanolamine, fatty acid (including long chain fatty acid elongation), and isoprenoid biosynthesis. Phospholipase B was previously identified as pathogenicity factors in *N. fowleri* (Barbour and Marciano-Cabral, 2001). Also identified was a Rieske cholesterol C7(8)-desaturase (Supplementary materials 2, Supplementary Figure 3), a protein involved in sterol production that is absent in mammals, thus potentially representing a drug target.

Recent work shows that *N. gruberi* trophozoites prefer to oxidize fatty acids to generate acetyl-CoA, rather than use glucose and amino acids as growth substrates (Bexkens et al., 2018). Several genes involved in metabolism of both lipids and carbohydrates are up-regulated in highly pathogenic *N. fowleri.* Of interest are those that may be involved in metabolizing the polyunsaturated long chain fatty acids that are abundant in the brain, such as long chain fatty acyl CoA synthetase and delta 6 fatty acid (linoleoyl CoA) desaturase-like protein. Consistent with possible shifting carbon source usage or increased growth rates, mitochondrial and energy conversion genes are up-regulated, such as ubiquinone biosynthesis genes, isocitrate dehydrogenase (TCA cycle), complex I and complex III genes (oxidative phosphorylation), and a mitochondrial ADP/ATP translocase. Eight genes involved in amino acid metabolism are also up-regulated.

Intriguingly, we identified several areas of *N. fowleri* metabolism that may impact the human host. Glutamate is found in high millimolar concentrations in the brain, and is thought to be the major excitatory neurotransmitter in the central nervous system (Danbolt, 2001; Featherstone and Shippy, 2008). Several genes that function in glutamate metabolism are up-regulated following mouse passage, including kynurenine-oxoglutarate transaminase, glutamate decarboxylase, glutamate dehydrogenase, and isocitrate dehydrogenase (Figure 3, Supplementary Table 9). Multiple neurotropic compounds are generated via these enzymes, such as kynurenic acid, GABA, and NH4^+^. Kynurenic acid in particular has been linked to neuropathological conditions in tick-borne encephalitis (Holtze et al., 2012).

**Figure 3.**
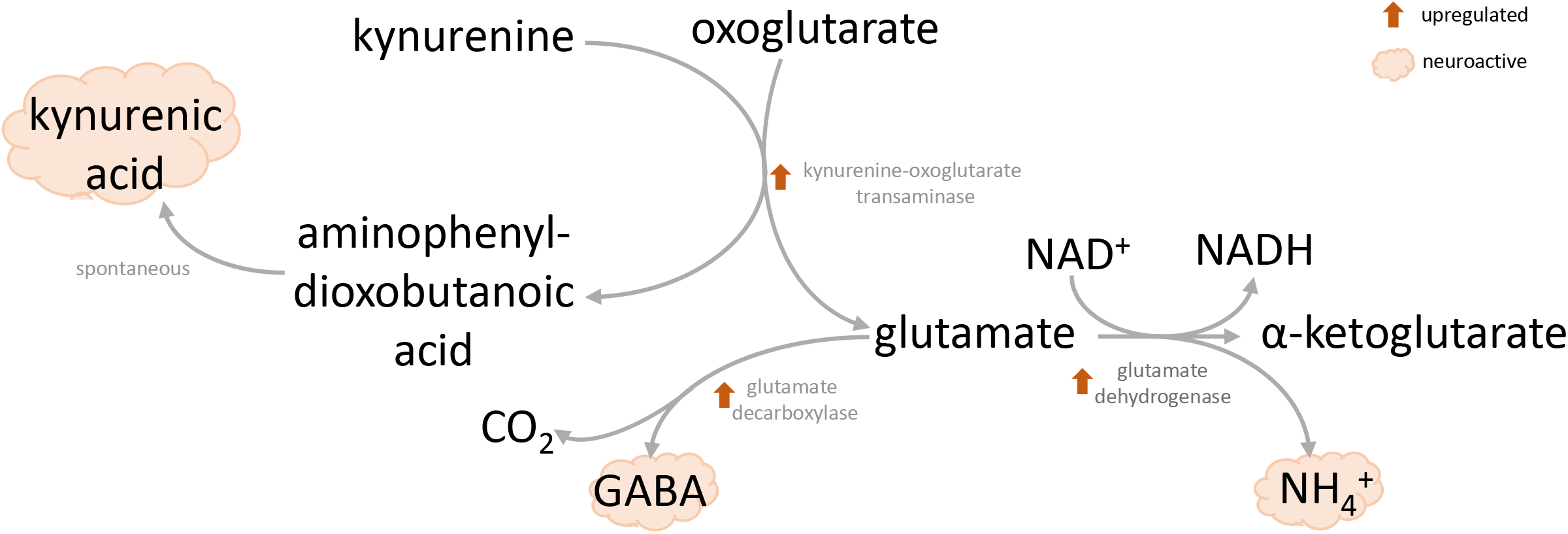
Highly pathogenic *N. fowleri* up-regulates enzymes producing neuroactive chemicals. Up-regulation of enzymes of glutamate metabolism following mouse passage of LEE *N. fowleri* suggests a strategy for ATP production *in vivo* and synthesis of neuroactive metabolites.

Ammonia transporters are also up-regulated (Supplementary Table 9), which may be one way to remove toxic ammonia from the cell, as *N. fowleri*, like *N. gruberi*, has an incomplete urea cycle (Opperdoes et al., 2011). Both glutamate and polyamine metabolism pathways discussed above generate NH4^+^as a by-product, and it is possible that this is secreted into the host brain and leads to pathological effects.

Finally, agmatine deiminase, which is involved in putrescine biosynthesis, is also up-regulated (Supplementary Table 9). This enzyme catalyzes the conversion of agmatine to carbamoyl putrescine, an upstream precursor to the polyamines glutathione and trypanothione (Colotti and Ilari, 2011). Trypanothione provides a major defense against oxidative stress, some heavy metals, and potentially xenobiotics in trypanosomatid organisms (e.g. *Leishmania*, *Trypanosoma*) (Fairlamb and Cerami, 1992), and has been isolated from *N. fowleri* trophozoites (Ondarza et al., 2006). Although a similar protective role for trypanothione has yet to be confirmed in *N. fowleri*, it is a critical component in trypanosomatid parasites (Krassner and Flory, 1971; Steiger and Steiger, 1977), and enzymes in this pathway may represent novel drug targets.

### Up-regulated genes with unclear roles in pathogenesis

There are many other genes that are up-regulated, but do not fall into one of the categories outlined above (Supplementary Table 9). Notably, one of these is a transcription factor of RWP-RK family, which has previously only been identified in plants and *D. discoideum,* functioning in plants to regulate responses in nitrogen availability, including differentiation and gametogenesis. While its role in amoebae is unclear, it may represent a potential drug target, as RWP-RK transcription factors are not present in human cells. Notably, of the 208 genes up-regulated in highly pathogenic *N. fowleri*, 10 are specific to *N. fowleri* (i.e. not found in other organisms), and 47 are found only in *N. fowleri* and *N. gruberi* (Supplementary Table 9). These represent unique potential targets against which anti-*Naegleria* therapeutics may be developed.

## Discussion

In this study, we provide the first genome-wide assessment of *N. fowleri* strain diversity and the first comprehensive systems-level analysis to understand why this species of *Naegleria* is a highly fatal human pathogen while other species are essentially benign.

Our work builds extensively upon previous understanding of *N. fowleri* pathogenicity. Our transcriptomic analysis revealed increased expression of several genes previously considered as pathogenicity factors (e.g., actin, the prosaposin precursor gene of Naegleriapore A and B, phospholipases and Nf314 (Cathepsin A) (Herbst et al., 2002; Hu et al., 1992; Sohn et al., 2010). In 2014, Zysset-Burri and colleagues published a proteomic screen of highly virulent versus weakly virulent *N. fowleri*, as a function of culturing cells with different types of media (Zysset-Burri et al., 2014). While there were clear differences between these strains, potentially due to genetic differences and the method of virulence induction, there were some shared pathways. This included villin and severin, which were both more abundant in high virulence *N. fowleri*, and are involved in actin cytoskeletal dynamics, as well as a phospholipase D homologue.

Our work, in combination with that of others, has allowed us to generate a model for pathogenicity in *N. fowleri* (Figure 4), hinging on both specific protein factors as well as whole cellular systems. Several previously identified pathogenicity factors are secreted proteases (e.g. metalloproteases, cysteine proteases, and pore-forming proteases) and phospholipases involved in host tissue destruction, which are likely to be secreted by the cell’s membrane trafficking system. Also part of this system – and a major source of differentially expressed genes – is the lysosomal degradation pathway. We identified many up-regulated cathepsin proteases, which function in the lysosome, and predict that they are involved in ingestion and breakdown of host material. Increased cellular ingestion goes hand in hand with cell growth and division, processes which we also see represented in the up-regulated gene dataset: genes involved in protein synthesis, metabolism, and mitochondrial function. This includes several metabolic pathways that produce compounds that could interact with the host immune system or have neurotropic effects. Finally, a major part of *N. fowleri*’s pathogenesis undoubtedly involves cell motility and phagocytosis, which are almost always actin-mediated processes. Because the cytoskeleton is central to *N. fowleri* viability and pathogenesis, and because actin-based processes likely dictate pathogenic behaviors, these analyses provide a solid framework for future investigation into *N. fowleri* virulence as well as potential drug targets.

**Figure 4.**
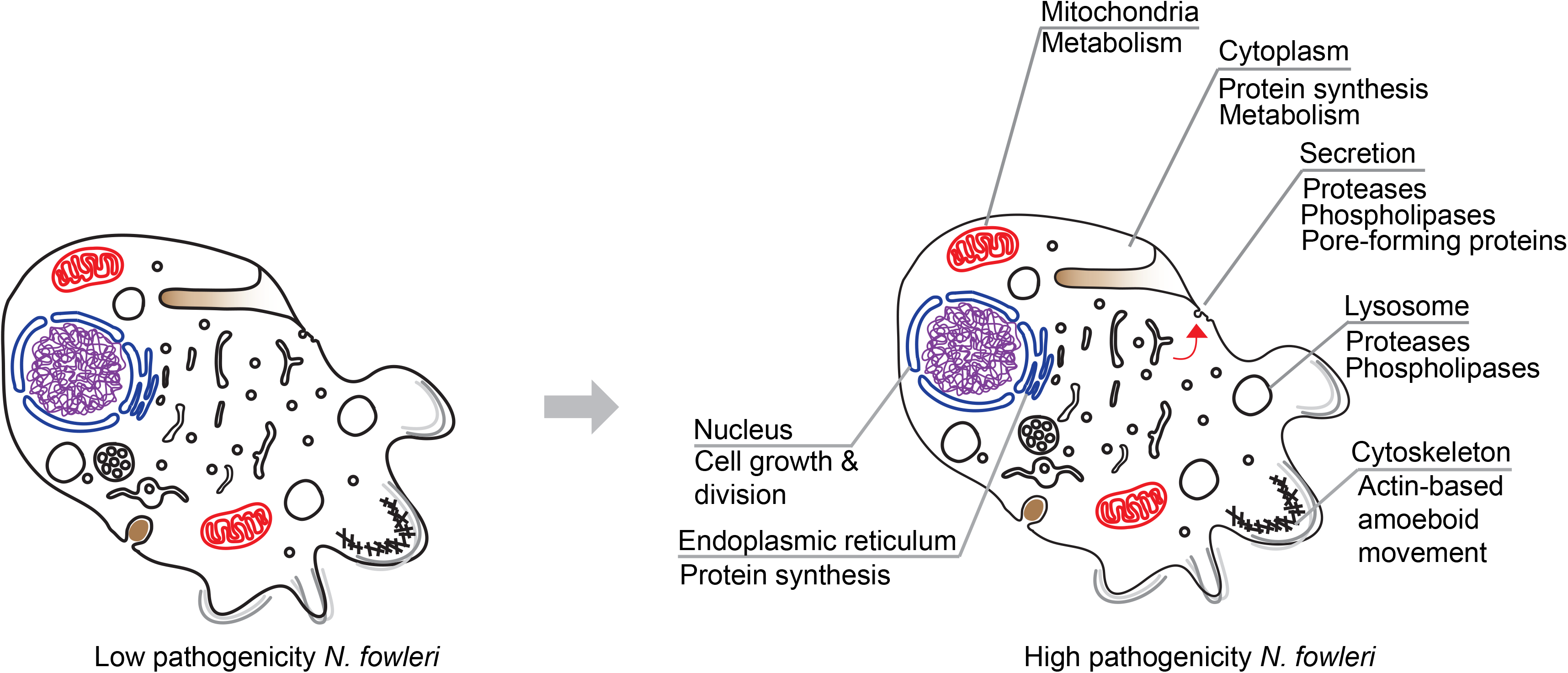
Model of *N. fowleri* pathogenicity. Aspects of cellular function that are likely relevant to *N. fowleri* pathogenicity are indicated on the cartoon of high pathogenicity *N. fowleri* (right), as compared with low pathogenicity *N. fowleri* (left). This model does not represent an exhaustive list of all identified pathogenicity factors, but rather maps the system-level changes in *N. fowleri* based on the results of our differential gene expression analysis.

Our comparative genomics and transcriptomics approaches have not only yielded the first systems level model of pathogenicity, but it has also identified many putative novel pathogenicity factors in *N. fowleri* (Figure 5). We identified key individual targets, such as the S81 protease and two cathepsin B proteases, which are both missing from *N. gruberi*, and differentially expressed in mouse passaged *N. fowleri*. Moreover, taking a hierarchical approach of overlapping criteria we can distill from high-throughput RNA-Seq data a catalogue of novel potential pathogenicity factors. Over 450 genes (458) are shared by *N. fowleri* strains to the exclusion of *N. gruberi*, while 315 are differentially expressed upon pathogenicity inducing conditions; both are logical criteria for their consideration as potential pathogenicity factors. Annotation as “unknown function” was taken as a criterion for novelty, but not necessarily pathogenicity. At the intersection of these criteria, there are 390 genes of unknown function that are specific to *N. fowleri*, and 115 genes that are specific to *N. fowleri* and are up-regulated following mouse-passage. Sixteen genes fulfill all three criteria; they are specific to *N. fowleri*, are up-regulated, and have no putative function. Notably, 90 of the up-regulated genes do not appear to have a human orthologue and represent potential novel drug targets. Genetic tools once developed in this lineage, can be used to functionally characterize the most promising candidate genes and better study *N. fowleri* cell biology, improving our understanding of why and how it is so virulent.

**Figure 5.**
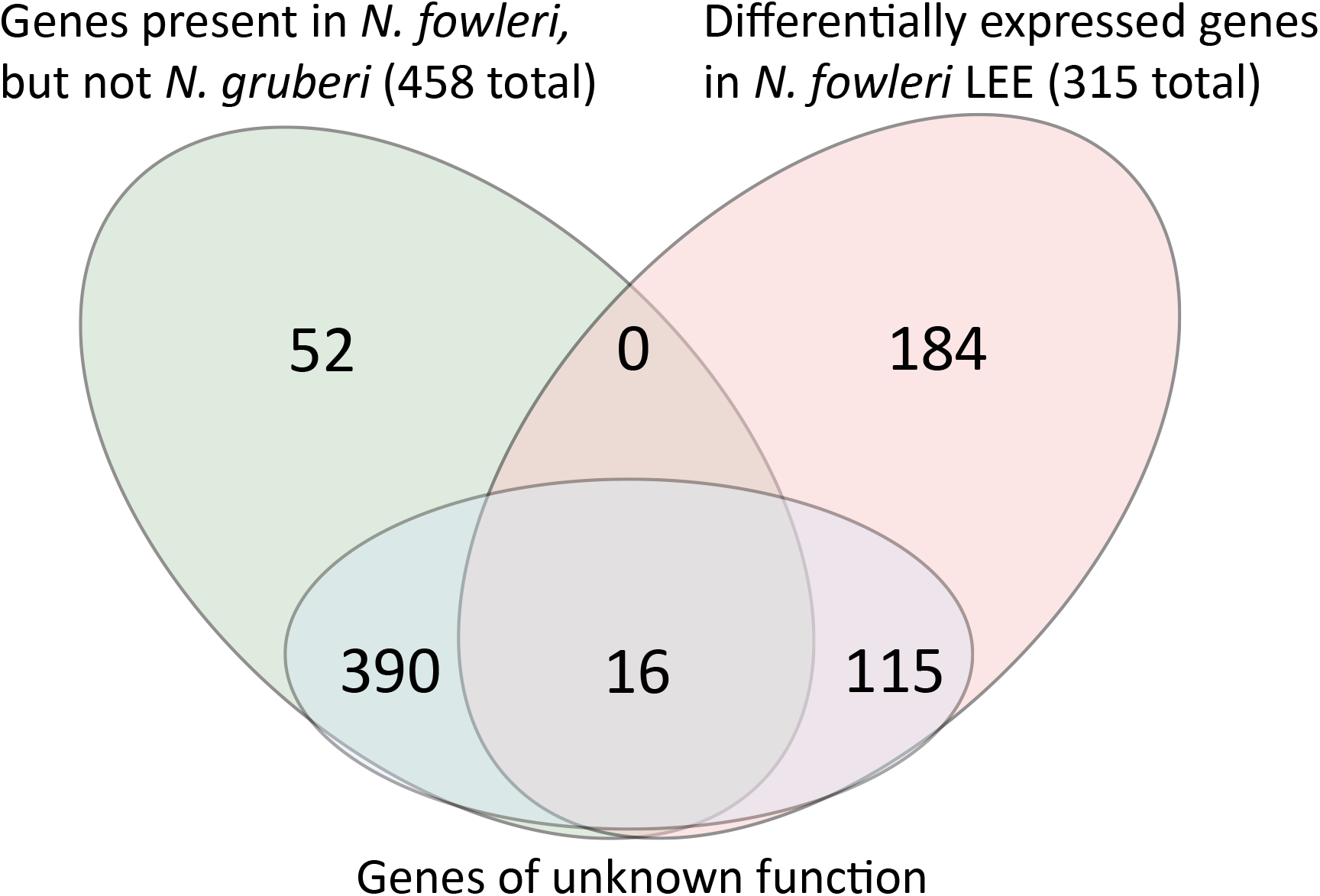
Venn diagram showing the overlap between differentially expressed genes in highly pathogenic *N. fowleri*, those which have no clear homologues in *N. gruberi*, and those of unknown function.

### Methods Summary

Genomic and transcriptomic data from three strains of *Naegleria fowler*i, including one strain passage axenically and through mice to compare DE genes implicated in pathogenicity, were obtained by Illumina sequencing. Gene models, expressions values, and comparative analyses were performed using a variety of computational tools. See Extended Methods for details (Supplementary Material 5).

## Acknowledgements

EKH was supported by an AHFMR Fulltime Graduate Studentship and a Vanier Canada Graduate Scholarship. JBD is the Canada Research Chair in Evolutionary Cell Biology. This work was also funded by a Royal Society International Exchanges grant (2015/R1-IE150049) jointly to ADT and JBD. Work in the Elias Lab is supported by: CePaViP (CZ.02.1.01/0.0/0.0/16_019/0000759) and Czech Science Foundation project 18-18699S. This work was supported by CSIRO Land and Water (Australia).

## Author Contributions

Generated unpublished primary data included in the manuscript: EKH, AG, HCM, MJM, DCZ-B, FMC. Provided curated analysis of an encoded genomic system and display items (includes supervision of trainee): EKH, IM-R, RV, MvdG, MLG, AT, KV, SRN, ME, CS, MW, LF-L, JBD. Edited or provided intellectual input on the manuscript: EKH, AG, MvdG, MLG, ADT, KV, SRN, GM, NM, ME, MW, LF-L, GJP, TW, CC, JBD. Initial conception of the project: FMC, GJP, MWi TW, NM, CC, JBD

---

**Supplementary Figure 1.** Dot plot showing the presence of meiosis genes in *Naegleria* sp. Meiosis gene in *N. gruberi* are based on Fritz-Laylin *et al.*, 2010. A filled circle indicates the presence of a gene, while an unfilled circle indicates that no clear homologue could be identified. Numbers within the circle indicate multiple paralogues. Like *N. gruberi*, *N. fowleri* encodes a highly complete complement of meiotic genes.

**Supplementary Figure 2.** Gene repertoire of the sterol synthesis pathway in *N. fowleri*. For comparison, the distribution of orthologues from *S. cerevisiae*, *H. sapiens* and *A.thaliana*, as well as the non-pathogenic *N. gruberi*, are shown. Double-dots indicate two orthologues performing one step (i.e. SMO1 and SMO2) or two orthologues performing similar reactions (i.e. SMT1 and SMT2). In the case of non-homologous families ERG2/EBP/HYD1 and ERG4/DHCR24/DWF1, the presence of one or the other copy is indicated by orange or blue dots as indicated in the gene names, whereas the presence of both copies is indicated by green dots. The Rieske cholesterol C7(8)-desaturase is shown apart because is not involved in the canonical sterol pathways.

**Supplementary Figure 3.** Partial alignment of cholesterol C7(8)-desaturases. Red boxes highlight the Rieske [2Fe-2S] motif (CXHX16CX2H) and the non-heme iron binding motif ((D/E)X3DX2HX4H). Asterisks indicate consensus residues important for catalysis.

**Supplementary Figure 4.** The *N. fowleri* genome encodes an extensive actin cytoskeletal repertoire. Formin family proteins typically nucleate and elongate actin filaments, moving progressively with the barbed end and recruiting profilin-bound actin monomers to the growing end of the filament. Another nucleator, the Arp2/3 complex, typically polymerizes a new filament from the side of a pre-existing filament following activation by a WASP-family protein such as WASP, SCAR/WAVE, and/or WASH. *N. fowleri* also encodes myosin motor proteins, including myosin I and II. Finally, *N. fowleri* encodes cofilin and members of the gelsolin/villin superfamily predicted to depolymerize actin filaments. In all, the *N. fowleri* genome encodes at least 22 actins, 14 formins, 11 myosins, 4 gelsolin/villin superfamily proteins, and 3 profilins, in addition to all the subunits of the Arp2/3 complex, and 3 WASP family proteins and their respective complexes.

**Supplementary Figure 5.** Phylogenetic analysis of Ras family genes in *Naegleria* species and other selected eukaryotes. Portrayed is a maximum likelihood tree (RAxML, LG+Γ model). Bootstrap support values were calculated using the rapid bootstrapping (Rboot) algorithm of the RAxML program. The robustness of the tree topology was also assessed by the IQ-tree with LG+F+G4 model (the model selected by the program itself) with the ultrafast bootstrap (UFboot) algorithm (1000 replicates) and the SH-aLRT test (1000 replicates). Circles at branches correspond to bootstrap values indicated in the legend. The bar on the top corresponds to the estimated number of substitutions per site. The identity of the *N. fowleri* genes is provided in Supplementary Table 7, sheet 2. Manually corrected gene models *; Newly created gene models**.

**Supplementary Figure 6.** Phylogenetic analysis of Rab family genes in *Naegleria* species and other eukaryotes from Elias et al., 2012 (Elias et al., 2012). Portrayed is a maximum likelihood tree (RAxML, LG+Γ model). Bootstrap support values were calculated using the rapid bootstrapping (Rboot) algorithm of the RAxML program. The robustness of the tree topology was assessed also by the IQ-tree with LG+G4 model (the model selected by the program itself) with the ultrafast bootstrap (UFboot) algorithm (1000 replicates) and the SH-aLRT test (1000 replicates). Circles at branches correspond to bootstrap values indicated in the legend. The bar on the top corresponds to the estimated number of substitutions per site. The identity of the *N. fowleri* genes is provided in Supplementary Table 7, sheet 2. Manually corrected gene models *; Newly created gene models**.

**Supplementary Figure 7.** Multidomain proteins with a Ras superfamily GTPase domain found in *Naegleria* species. Selected proteins containing more than one domain are schematically depicted here. The positions of domains are according to NCBI's conserved domain database, except the C-terminal part of Gpa18 protein which was identified by manual inspection of the multiple sequence alignment of *Naegleria spp.* G*α* proteins, as this fragment was too small to be detected by NCBI’s conserved domain search. Ras superfamily domains are depicted in orange with a label specifying a subgroup of the Ras superfamily. ZnF UBP = Ubiquitin carboxyl-terminal hydrolase-like zinc finger; ANKs = multiple ankyrin repeats; G – alpha (G*α*) = G protein alpha subunit; STKc = serine/threonine protein kinases. Nfo: *N. fowleri*; Ngr: *N. gruberi*; *: manually corrected gene model. The scale bar shows the length of 100 amino acid residues.

**Supplementary Figure 8.** MA plot and Volcano plot of genes differentially expressed in highly pathogenic *N. fowleri* LEE strain. MA plots show the log fold change of each gene relative to the log of the mapped read counts, while volcano plots scale the False Discovery Rate (FDR) to log fold change. Each gene is represented by a dot, and red dots indicate those which meet the differential expression criteria.

**Supplementary Figure 9.** Protease gene abundance in *Naegleria* per MEROPS protease family. For each MEROPS protease family, the number of predicted proteins in each strain is shown. The S81 family (boxed) is the only family without a homologue in *N. gruberi*. The only other known S81 family protease is a destabilase protein in *Hirudo medicinalis*, the European medicinal leech. Otherwise, most protease families could be identified in all four genomes, with similar numbers of homologues. The exception is the C01 cysteine protease family, which is analyzed in Supplementary Figure 10.

**Supplementary Figure 10.** Phylogenetic analysis of the C01 cysteine protease subfamily in *N. fowleri* and *N. gruberi*. Node values are listed as Phylobayes/RAxML (posterior probability/bootstrap), and as symbols indicating a minimum level of support as shown in the inset. Node values are shown on the best Bayesian topology. Sequences with signal peptides have red text, those with potential signal peptides (score near cutoff) have purple text, and those without identifiable signal peptides have blue text. Asterisks (*) indicate genes that are up-regulated in highly pathogenic *N. fowleri*.

