## Supplementary Figure 1 for "A comparative ‘omics approach to candidate pathogenicity factor discovery in the brain-eating amoeba *Naegleria fowleri*"

| | *N. gruberi* | *N. fowleri* V212 | *N. fowleri* 30863 | *N. fowleri* 986 |
|---|---|---|---|---|
| Spo11* | ❷ | ❷ | ❷ | ● |
| Mre11 | ❷ | ● | ● | ● |
| Rad50 | ● | ● | ● | ● |
| Rad1 | ● | ● | ● | ● |
| ERCC4 | ● | ● | ● | ● |
| Hop1* | ● | ● | ● | ● |
| Hop2* | ❷ | ❷ | ❷ | ❷ |
| Mnd1* | ● | ● | ● | ● |
| Rad52 | ❸ | ❷ | ❷ | ❷ |
| Dmc1* | ● | ● | ● | ● |
| Rad51 | ❸ | ❺ | ❺ | ❺ |
| Msh2 | ● | ● | ❷ | ● |
| Msh6 | ❷ | ● | ● | ● |
| Msh3 | ● | ● | ● | ● |
| Msh4* | ● | ● | ● | ● |
| Msh5* | ● | ● | ● | ● |
| Mlh1 | ● | ● | ● | ● |
| Mlh2 | ○ | ○ | ○ | ○ |
| Mlh3 | ● | ● | ● | ● |
| Pms1/2 | ❷ | ❷ | ❷ | ❷ |
| Mer3* | ● | ● | ● | ● |
| Smc1 | ● | ● | ● | ● |
| Smc2 | ● | ● | ● | ● |
| Smc3 | ● | ● | ● | ● |
| Smc5 | ● | ● | ● | ● |
| Rad18 | ● | ● | ● | ● |
| Rad21 | ● | ● | ● | ● |
| Rec8* | ○ | ○ | ○ | ○ |
| Pds5 | ● | ● | ● | ● |
| Sec3 | ● | ● | ● | ● |

Based on Malik *et al.* 2008 and Fritz-Laylin *et al.* 2010
* Considered to be meiosis-specific in Malik *et al.* 2008
