## Supplementary Figure 2 for "A comparative ‘omics approach to candidate pathogenicity factor discovery in the brain-eating amoeba *Naegleria fowleri*"

| | *S. cerevisiae* | *H. sapiens* | *A. thaliana* | *N. gruberi* | ***N. fowleri*** |
|---|---|---|---|---|---|
| **Squalene monooxygenase (ERG1/SQLE/SQE)** | ● | ● | ● | ● | ● |
| **Oxidosqualene cyclase (ERG7/LSS/CAS1)** | ● | ● | ● | ● | ● |
| **C-14 demethylase (ERG11/CYP51A1/CYP51G1)** | ● | ● | ● | ● | ● |
| **C14-reductase (ERG24/TM7SF2/FK)** | ● | ● | ● | ● | ● |
| **C-4 methyl oxydase (ERG25/SC4MOL/SMO1-2)** | ● | ● | ●● | ●● | ●● |
| **C-3 dehydrogenase (ERG26/NSDHL/AT3betaHSD)** | ● | ● | ● | ● | ● |
| **3-keto reductase (ERG27/HSD17B7/?)** | ● | ● | ○ | ○ | ○ |
| **ER anchor (ERG28/C14orf1/ERG28)** | ● | ● | ● | ● | ● |
| **C-24 methyltransferase (ERG6/ - /SMT1-2)** | ● | | ●● | ● | ● |
| **Δ8-Δ7 isomerase (ERG2/EBP/HYD1)** | ● (orange) | ● (blue) | ● (blue) | ● (green) | ● (green) |
| **C-5 desaturase (ERG3/SC5DL/STE1)** | ● | ● | ● | ● | ● |
| **Δ7(8)-reductase (- /DHCR7/DWF5)** | | ● | ● | ● | ● |
| **C-22 desaturase (ERG5/ - /CYP710A1)** | ● | | ● | | |
| **Δ24-reductase (ERG4/DHCR24/DWF1)** | ● (orange) | ● (blue) | ● (blue) | ● (orange) | ● (orange) |
| **Cyclopropyl isomerase (CPI1)** | | | ● | ● | ● |
| **Cholesterol C7(8)-desaturase (Nvd/DAF36/Des7)** | | | | ● | ● |
