## Supplementary Material 1 for "A comparative ‘omics approach to candidate pathogenicity factor discovery in the brain-eating amoeba *Naegleria fowleri*"

*Transcription factor identification*

We identified likely transcription factors (TFs) by scanning the amino acid sequences of predicted protein coding genes for putative DNA binding domains (DBDs) using the procedures described in Weirauch *et al.* ^1^. Briefly, we scanned all protein sequences for putative DBDs using the 81 Pfam^2^ models listed in Weirauch and Hughes^3^ and the HMMER tool^4^, with the recommended detection thresholds of Per-sequence Eval < 0.01 and Per-domain conditional Eval < 0.01. Each protein was classified into a family based on its DBDs and their order in the protein sequence (e.g., bZIPx1, AP2x2, Homeodomain+Pou). Using the above procedure, we identified a total of 203, 196, and 196 putative TFs in the genomes of *N. fowleri* strains 30863, 986, and V212, respectively. These values are similar to those reported previously for *N. gruberi* (243), especially when accounting for the fact that *N. gruberi* has ~30% more genes than *N.fowleri*. A few TF families are under-represented in *N. fowleri* relative to *N. gruberi* (e.g., zinc clusters, AT hooks, and HMG boxes), and one family (GATA) is substantially expanded (10 vs 4 genes).

1.	Weirauch, M. T. *et al.* Determination and inference of eukaryotic transcription factor sequence specificity. *Cell* **158**, 1431–1443 (2014).

2.	Finn, R. D. *et al.* The Pfam protein families database. *Nucleic Acids Res.* **Database I**, D211-22 (2010).

3.	Weirauch, M. T. & Hughes, T. R. A catalogue of eukaryotic transcription factor types, their evolutionary origin, and species distribution. *Subcell. Biochem.* **52**, 25–73 (2011).

4.	EDDY, S. R. A NEW GENERATION OF HOMOLOGY SEARCH TOOLS BASED ON PROBABILISTIC INFERENCE. in *Genome Informatics 2009* (2009). doi:10.1142/9781848165632_0019
