## Supplementary Material 2 for "A comparative ‘omics approach to candidate pathogenicity factor discovery in the brain-eating amoeba *Naegleria fowleri*"

**Sterols metabolism genes from *Naegleria fowleri***

Sterols are fundamental lipids in eukaryotes. They are important components of the plasma membrane, playing relevant structural roles and also participating in cellular signaling. Due to its essentiality and prevalence in eukaryotic microorganisms (Desmond and Gribaldo, 2009), sterol metabolism is a frequent therapeutic target against pathogenic fungi and microbial parasites including kinetoplastids and various amoebae (Choi et al., 2014).

A sterol biosynthesis pathway was proposed two decades ago in the non-pathogenic *Naegleria gruberi* and *N. lovaniensis* (Raederstorff and Rohmer, 1987). Like fungi and kinetoplastids, *Naegleria* species use ergosterol as the major sterol component of their membranes. While fungi and kinetoplastids use lanosterol as a starting point, *Naegleria* species produce ergosterol through cycloartenol as the first cyclic product of the pathway (a characteristic shared with some photosynthetic lineages). Because efforts in developing effective treatments for *N. fowleri* infections rely on repurposing of existing drugs (e.g. antifungals), it is imperative to determine the identity of the enzymatic steps involved in the biosynthesis of ergosterol in *N. fowleri* (Debnath et al., 2017).

The putative sterol biosynthesis pathway of *N. fowleri* was reconstructed using BLASTp to mine the predicted proteomes from the three strains studied here. Amino acid sequences of sterol metabolism proteins from *Homo sapiens*, *Saccharomyces cerevisiae* and *Arabidopsis thaliana*, were used as queries. We have also searched for orthologues of the cholesterol C7(8)-desaturase (EC 1.14.19.21).

We have found the complete set of proteins for the production of sterols in the three *N. fowleri* strains investigated (Supplementary Figure 2). We have also detected putative orthologues of the Rieske cholesterol C7(8)-desaturase (Supplementary Figure 3). This enzyme converts cholesterol into 7-dehydrocholesterol during steroid hormones synthesis in ecdysozoans (Wollam et al., 2011; Yoshiyama-Yanagawa et al., 2011), in the ciliate *Tetrahymena thermophila* (Najle et al., 2013), and was also proposed to be involved in the pathway of conversion of diet cholesterol into ergosterol in the unicellular amoeba *Capsaspora owczarzaki* (Najle et al., 2016). Importantly, cholesterol 7(8)-desaturase is highly conserved in animals except mammals (Najle et al., 2013), thus providing a potential candidate target to design drugs to abolish or restrict the synthesis of sterols in *N. fowleri* without affecting the human host.

Like in *N. gruberi*, there are no orthologs of sterol C7-reductase in *N. fowleri*. This is not surprising if we assume that the pathway will lead to ergosterol as the end product, like in the non-pathogenic *Naegleria* species. Also, we were unable to detect an ortholog of ERG5/CYP710A1, the cytochrome P450 C22-desaturase. Desaturation at position C22(23) of the lateral chain is necessary for the production of ergosterol. However, the absence of an ERG5/CYP701A1 ortholog does not necessarily mean that *Naegleria* *spp*. are not capable of performing this enzymatic activity. The choanoflagellate *Monosiga brevicolis* (Kodner et al., 2008) and the ciliate *T. thermophila* (Najle et al., 2019) are examples of species known to produce delta-22-sterols, without having a canonical P450 sterol C22-desaturase (Fu et al., 2009).

None of the genes encoding sterol metabolism enzymes exhibited significant changes in transcript levels in our RNAseq experiments. The only gene related to this metabolic pathway that is differentially expressed is the squalene synthase (SQS). However, squalene is the building block of many isoprenoids, not only sterols. Thus, a direct link between the overexpression of SQS and the overproduction of sterols (and hence its involvement in the infection process) cannot be suggested.
