## Supplementary Material 3 for "A comparative ‘omics approach to candidate pathogenicity factor discovery in the brain-eating amoeba *Naegleria fowleri*"

*Analysis of the cytoskeletal protein complement in* N. fowleri

The actin and microtubule cytoskeletons coordinate and execute nearly every cellular function, including migration, endocytosis, and cell division. In addition to clear roles in cell growth and viability, these functions likely drive pathogenesis (reviewed in (Siddiqui et al., 2016)); after inhalation of contaminated water, *N. fowleri*  migrates to and within the brain (Rojas-Hernandez et al., 2004), where endocytosis of host material (Brown, 1978; Visvesvara and Callaway, 1974) and release of mucus- and tissue-degrading enzymes cause damage and inflammation (Aldape et al., 1994; Martinez-Castillo et al., 2017; Serrano-Luna et al., 2007). To understand the mechanisms underlying these pathogenic behaviors, we need a clear picture of the composition of the microtubule and actin cytoskeletons.

Much of what we know about *Naegleria*’s cytoskeleton pertains to its unique differentiation from amoebae into flagellates. During this transition, *Naegleria* synthesize all the proteins required to build flagella, including basal bodies and flagellar tubulin (Fritz-Laylin et al., 2010a; Fritz-Laylin and Cande, 2010; Lai et al., 1979; Patterson et al., 1981). In addition, *Naegleria* presumably uses microtubules for chromosome segregation during closed mitosis (Gonzalez-Robles et al., 2009; Walsh, 2012).  We find that *N. fowleri* and the nonpathogenic species *Naegleria gruberi* each possess an extensive repertoire of tubulins and microtubule associated proteins (Supplementary Table 5). We identified over 140 genes encoding proteins involved in the microtubule cytoskeleton in *N. fowleri*, including; several tubulins (including >13 alpha and beta tubulins, gamma, delta, and epsilon tubulins), motor proteins (>30 kinesins and 6 dyneins (based on the number of heavy chains)), as well as proteins associated with interflagellar transport, basal body assembly and structure (Fritz-Laylin et al., 2010a; Fritz-Laylin and Cande, 2010), and many other microtubule binding proteins.

Despite this extensive microtubule gene repertoire, infectious *Naegleria* amoebae move, eat, and divide without cytoplasmic microtubules (Gonzalez-Robles et al., 2009; Rojas-Hernandez et al., 2004; Walsh, 1984; Walsh, 2007). Moreover, previous studies have suggested that actin, actin binding proteins, or upstream actin regulators may correlate with virulence (Jamerson et al., 2017; Zysset-Burri et al., 2014). Therefore, a thorough understanding of *Naegleria*’s control of actin dynamics remains critical to understanding its pathogenesis.

In cells, actin polymer formation requires proteins called nucleators (Campellone and Welch, 2010; Dominguez, 2016). The Arp2/3 complex is a nucleator that typically generates branched actin networks that are useful for cell motility and vesicle trafficking (Campellone and Welch, 2010; Rotty et al., 2013). We predict that *N. fowleri* can form these branched actin networks, as we identified all seven subunits of the Arp2/3 complex (Supplementary Table 5). We also found upstream Arp2/3 activators from the WASP-family, including WASP and all components of the WASH and SCAR/WAVE complexes. The presence of WASP and SCAR/WAVE together is indicative of the capacity for pseudopod-based alpha-motility (Fritz-Laylin et al., 2017), which is consistent with microscopic observations (Sohn et al., 2010; Walsh, 2007).

In addition to Arp2/3 complex-mediated nucleation, cells also use formin family proteins to nucleate and elongate actin polymers (Supplementary Figure 4). Formin proteins have highly variable domain architectures, and can also bundle and sever actin filaments (Breitsprecher and Goode, 2013). We identified 14 formin homology 2 (FH2) domain-containing proteins, and of those 12 have similar domains and organization to diaphanous related formins (DRFs), and two contain phosphatase and tensin (PTEN) domains. Because amoebae rely on actin for their cytoskeleton, and because actin regulation often occurs at the nucleation level, these Arp2/3- and formin-mediated pathways of actin polymerization likely drive most cellular functions.

As an alternative strategy to identify actin related pathogenicity factors, we compared *N. fowleri* cytoskeletal genes to those of its nonpathogenic relative, *N. gruberi* (Supplementary Table 5). Generally, *N. fowleri* and *N. gruberi* harbor similar numbers of the same genes. Because actin is regulated largely at the level of nucleation, we classified the *N. gruberi* and *N. fowleri* formin families by domain composition and found a single difference in one formin conserved in the two species. In this formin (62754), the *N. fowleri* homolog has a putative lipid-binding PTEN domain that is missing from that of *N. gruberi.* While the comparison between *Naegleria* strains did not reveal any major differences in the cytoskeletal repertoires that can immediately explain the differences in pathogenicity, it is notable that humans do not encode formins of the PTEN family. Therefore, if these PTEN formins are responsible for any vital *Naegleria* processes, they may represent useful drug targets.

Because the cytoskeleton is central to *N. fowleri* viability and pathogenesis, and because actin-based processes likely dictate pathogenic behaviors, these analyses provide a solid framework for future investigation into *N. fowleri* virulence as well as potential drug targets.
