## Supplementary Material 4 for "A comparative ‘omics approach to candidate pathogenicity factor discovery in the brain-eating amoeba *Naegleria fowleri*"

**Ras superfamily GTPases in *Naegleria* spp.**

The Ras superfamily of GTPases represents a vast group of proteins involved in endomembrane transport, cell signalling, actin dynamics, cilium-associated functions, and many other eukaryote-specific processes (Wennerberg et al. 2005). The previously published *Naegleria gruberi* genome sequence revealed a plethora of Ras superfamily members, including “monomeric Ras-like GTPases” (182 genes identified by the Pfam profile PF00071) and alpha subunits of heterotrimeric G proteins (Gα; 39 genes identified by the Pfam profile PF00503) (Fritz-Laylin et al. 2010). In order to assess the differences between *N. gruberi* and *N. fowleri*, we annotated the three strains of *N. fowleri* and also carefully reannotated the complement of Ras superfamily genes in the *N. gruberi* genome.

Reannotation of the complement of Ras superfamily genes in *N. gruberi* yielded over 350 Ras superfamily genes. Compared to the original annotation (Fritz-Laylin et al. 2010), many incorrectly predicted gene models were fixed and a number of genes that had escaped annotation were identified and annotated anew (the reannotated set of the *N. gruberi* Ras superfamily GTPases is listed in Supplementary Table 7, sheet 1). The Ras superfamily complement in *N. fowleri* is much less expanded, amounting to “only” over 200 genes (all listed in Supplementary Table 7, sheet 2), which is nevertheless still a lot compared to the numbers usually seen in eukaryotes. Little, if any, differences were found between the three *N. fowleri* strains (Supplementary Table 7, sheet 2). A substantial part (>180) of the genes exhibits an obvious one-to-one orthology relationship between *N. gruberi* and *N. fowleri* (Supplementary Table 7, sheet 2), but paralog expansions have extensively shaped the complements of Ras superfamily genes after the divergence of the lineages leading these two species. However, *N. gruberi* and much less frequently *N. fowleri* exhibit Ras superfamily genes lacking obvious orthologs in the other species. These may represent extremely divergent lineage-specific paralogs or genes predating the two species but lost in one or the other species. Genome analyses of other heteroloboseans are required to distinguish between these two possibilities.

Our analyses revealed that *Naegleria* inherited a large complement of Ras superfamily genes form an ancestor shared with other eukaryotes: both *Naegleria* spp. exhibit apparent orthologs of more than 50 Ras superfamily members from eukaryotes outside Heterolobosea (Supplementary Table 7, sheet 2; note that eukaryote-wide phylogenetic history of some subgroups of the Ras superfamily, e.g. the Gα proteins, has not been extensively investigated yet, so our result concerning orthologous relationships between *Naegleria* and other eukaryotes is a minimal estimate). There are few known conserved Ras superfamily members common in other eukaryotes yet missing in *Naegleria*, notable examples being Rab24 (Elias et al. 2012) or proteins of the Roco family (Marín et al. 2008). It was previously reported that *N. gruberi* has the GTPase Rheb (van Dam et al. 2011a), but the respective gene (XP_002683011.1) and its *N. fowleri* ortholog (Ras30) are more likely divergent paralogs of the true Ras (Supplementary Figure 3), whereas the *bona fide* Rheb was lost in the *Naegleria* lineage, yet retained by some other heteroloboseans (Záhonová et al. 2018).

Interesting commonalities as well as differences are seen when individual main subgroups of the Ras superfamily are analysed in the two *Naegleria* species. 44 Rab and Rab-like genes were previously reported from *N. gruberi* (Elias et al. 2012). We now additionally identified a number of more divergent Rab-like genes in the *N. gruberi* genome, expanding the size of the Rab(-like) family to nearly 90 loci in this species, whereas the equivalent group of genes comprises at least 55 Rab-like loci in *N. fowleri*. The difference stems essentially only from differential expansion of three subgroups of divergent Rab-like genes lacking discernible close relatives in eukaryotes outside *Naegleria* (Supplementary Figure 4). Interestingly, some members of the three divergent Rab-like clades in both *Naegleria* species seem to be affected by mutations disrupting the coding sequence as predicted based on conservation with the related genes, suggesting a recent dynamic birth-and-death evolution. Proteins in each of the three subgroups possess a conserved N-terminal extension upstream of the GTPase domain, in at least some of these proteins identifiable as the F-box domain (see also Supplementary Figure 5). One of these three subgroups has a conserved C-terminal prenylation motif (in the CaaX form; Leung et al. (2006)), suggesting that it is geranylgeranylated (or farnesylated) and attaches to membranes like standard Rabs and some other Ras superfamily GTPases. No specific functional prediction for these proteins is possible from the sequence analysis only, but their involvement in processes at the organism-environment interface seems likely.

We detected 80 and more than 40 genes representing the Ras family in *N. gruberi* and *N. fowleri*, respectively. These include the previously detected CPRas protein characterized by a unique circularly-permuted GTPase domain (Elias & Novotny, 2008) and three paralogs of the GTPase Rap1 (Supplementary Figure 3). The remaining Ras family genes in *Naegleria* species all seem to derive from the ancestral true Ras protein (van Dam et al. 2011b) based on sequence similarity comparisons, although many of the paralogs in *Naegleria* spp. are too divergent to prove this by phylogenetic analyses. The Rho family is represented by nearly 50 and more than 80 genes in *N. fowleri* and *N. gruberi*, respectively. Interestingly, the family also includes the RhoBTB type (proteins with a Rho-like GTPase domain fused to a tandem of BTB domains) so far known only from metazoans and the amoebozoan *Dictysotelium* (Ji & Rivero, 2016). Expanded families of Ran GTPase homologs are rarely seen in eukaryotes, so it is interesting to note that *N. fowleri* and *N. gruberi* harbor six and seven Ran or Ran-like genes, respectively (including one presumable pseudogene in the former, Supplementary Table 7). These include not only paralogs highly similar to canonical Ran from other eukaryotes and thus apparently mediating nucleocytoplasmic transport (Görlich & Mattaj, 1996), but also several unusual more divergent paralogs, some of them provided with N-terminal extra domains (see below and Supplementary Figure 5), suggesting their possible recruitment for novel cellular roles. The Arf/Arl/Sar1 family is not particularly expanded in *Naegleria*, but the two species analysed do differ in a complement of several divergent taxon-specific paralogs of unknown function. Finally, our reanalysis of the *N. gruberi* genome revealed that the Gα family (alpha subunits of heterotrimeric G-proteins) is even larger than reported previously (39 genes identified by Fritz-Laylin et al. 2010). We now detected 60 Gα genes in this species, whereas *N. fowleri* has a smaller, but still unusually large complement of >30 Gα genes.

The GTPase domain of the Ras superfamily typically exists as a stand-alone protein (i.e. a “small GTPase”), possibly with an (often prenylated or acylated) N- or C-terminal extension mediating membrane attachment, but it can be also incorporated into larger proteins featuring additional functional domains. In addition to several multi-domain Ras superfamily proteins representing ancestral or at least widespread forms, e.g. CPRas, RhoBTB, Miro (Elias & Novotny, 2008; Vlahou et al. 2011; Ji & Rivero, 2016), both *Naegleria* spp. exhibit a number of genes with novel domain architectures including a Ras superfamily GTPase domain as one of the elements (listed in Supplementary Table 7, examples provided in Supplementary Figure 5). The most frequent additional domain is the F-box found in combination with GTPase domains belonging to the Rab, Rho, Ran, and Gα families. Curiously, in the latter case the F-box domain is nested within the GTPase domain. We also identified one Ras protein and several Arf and Arf-like proteins combined with the BTB domain. Both F-box and BTB domains are implicated in protein-protein interactions, particularly in the context of protein ubiquitination as components of ubiquitin ligases (Kipreos & Pagano, 2000; Perez-Torrado et al. 2006), pointing to GTPase-dependent ubiquitination as a possibly significant way of cellular regulation in *Naegleria*. In addition to domains mentioned above, we also identified combinations of a Ras superfamily GTPase domain with, e.g., a serine/threonine protein kinase domain, LRR repeats, Ankyrin repeats, or Kelch motifs .

Our analyses revealed that amoeboflagellates of the genus *Naegleria* possess a highly complex and evolutionarily dynamic set of Ras superfamily genes that points to a hidden level of differentiation in cellular physiology of different *Naegleria* species. Interestingly, a similar pattern is seen in members of Amoebozoa (see, e.g., Sucgang et al. 2011), so we speculate that an amoeboid lifestyle generally requires a complex GTPase-based regulation of cellular activities. For example, in analogy to results obtained by studying metazoan and *Dictyostelium* amoeboid cells (Liu et al. 2016), proteins from the highly expanded Ras, Rho, and Gα families may underpin sophisticated functional networks responsible for sensing and reacting to various chemoattractants. Unfortunately, testing this hypothesis is precluded by the current lack of tools for genetic manipulation of *Naegleria* spp.

**Material and Methods**

We used genomic and predicted protein sequences of *N. gruberi* (Fritz-Laylin et al. 2010) and *N. fowleri* strain V212. Two additional *N. fowleri* strains (986 and 30863) were also analysed. For comparative analyses we also identified Ras superfamily genes from a set of reference, phylogenetically diverse species using data available in the NCBI database (http://www.ncbi.nlm.nih.gov). Ras superfamily gene sequences were detected and preliminarily identified using the program BLAST and its variants (Altschul et al. 1997). The protein sequences were manually inspected and prediction of the underlying gene models corrected whenever necessary. Multiple protein alignments were constructed using the program MAFFT (Katoh & Standley, 2013). All alignments were checked and if necessary, further edited manually using BioEdit (Hall 1999). Phylogenetic trees were computed using the maximum likelihood method implemented in the program RAxML-HPC with the LG+Γ model and branch support assessed by the rapid bootstrapping algorithm (Stamatakis, 2014) at the CIPRES Science Gateway portal (Miller et al. 2010). The robustness was additionally assessed by the maximum likelihood method implemented in the IQ-tree program (Kalyaanamoorthy et al. 2017) with branch supports assessed by the ultrafast bootstrap approximation (Minh et al. 2013) and SH-aLRT test (Guindon et al. 2010). Genes with one-to-one orthology relationships were detected by maximum likelihood phylogenetic analysis (IQ-tree) or BLASTp searches. Two genes were considered as one-to-one orthologs if they form a monophyletic group to the exclusion of other analysed sequences with support at least 80/95 (SH-aLRT support (%) / ultrafast bootstrap support (%)), or if they represent reciprocally best hits in BLASTp searches in our in-house Ras superfamily GTPase database containing also Ras superfamily genes from various other taxa. For the analysis of Rab proteins, the dataset from Elias et al. (2012) was used and expanded with sequences from *Naegleria* spp. Protein domains were searched using SMART (Letunic et al. 2015), Pfam (Finn et al. 2016) and NCBI's conserved domain database (Marchler-Bauer et al. 2015). Several suspicious domains detected only by NCBI's conserved domain with low e-value and/or representing only a small part of detected domain were excluded from the results.

**SuppFig5_RasGTPases.** Phylogenetic analysis of Ras family genes in *Naegleria* species and other selected eukaryotes. Portrayed is a maximum likelihood tree (RAxML, LG+Γ model). Bootstrap support values were calculated using the rapid bootstrapping (Rboot) algorithm of the RAxML program. The robustness of the tree topology was assessed also by the IQ-tree with LG+F+G4 model (the model selected by the program itself) with the ultrafast bootstrap (UFboot) algorithm (1000 replicates) and the SH-aLRT test (1000 replicates). Circles at branches correspond to bootstrap values indicated in the legend. The bar on the top corresponds to the estimated number of substitutions per site. The identity of the *N. fowleri* genes is provided in Supplementary Table 7, sheet 2. Manually corrected gene models *; Newly created gene models**.

**SuppFig6_RabGTPases.** Phylogenetic analysis of Rab family genes in *Naegleria* species and other eukaryotes from Elias et al., 2012. Portrayed is a maximum likelihood tree (RAxML, LG+Γ model). Bootstrap support values were calculated using the rapid bootstrapping (Rboot) algorithm of the RAxML program. The robustness of the tree topology was assessed also by the IQ-tree with LG+G4 model (the model selected by the program itself) with the ultrafast bootstrap (UFboot) algorithm (1000 replicates) and the SH-aLRT test (1000 replicates). Circles at branches correspond to bootstrap values indicated in the legend. The bar on the top corresponds to the estimated number of substitutions per site. The identity of the *N. fowleri* genes is provided in Supplementary Table 7, sheet 2. Manually corrected gene models *; Newly created gene models**.

**SuppFig7_MultidomainGTPases.** Multidomain proteins with a Ras superfamily GTPase domain found in *Naegleria* species. Selected proteins containing more than one domain are schematically depicted here. The positions of domains are according to NCBI's conserved domain database, except the C-terminal part of Gpa18 protein which was identified by manual inspection of the multiple sequence alignment of *Naegleria* spp. Gα proteins, as this fragment was too small to be detected by NCBI’s conserved domain search. Ras superfamily domains are depicted in orange with a label specifying a subgroup of the Ras superfamily. ZnF UBP = Ubiquitin carboxyl-terminal hydrolase-like zinc finger; ANKs = multiple ankyrin repeats; G – alpha (Gα) = G protein alpha subunit; STKc = serine/threonine protein kinases. Nfo: *N. fowleri*; Ngr: *N. gruberi*; *: manually corrected gene model. The scale bar shows the length of 100 amino acid residues.
