## Supplementary Material 5 for "A comparative ‘omics approach to candidate pathogenicity factor discovery in the brain-eating amoeba *Naegleria fowleri*"

*Culturing*

The Australian *Naegleria fowleri* isolate 986 was obtained from an operational drinking water distribution system in rural Western Australia and verified as *N. fowleri* using qPCR-melt curve analysis (Puzon et al., 2009) and DNA sequencing to confirm as Type 5 variant (De Jonckheere, 2011). The Australian *N. fowleri* type 5 was cultured on axenic media, modified Nelson’s Medium consisting of: 1 g/L Oxiod Liver Digest (Oxiod), 1 g/L Glucose, 24.0 g/L NaCl, 0.40 g/L MgSO4-7H2O, 0.402 g/L CaCl-2H2O, 14.2 g/L Na2HPO4, 13.60 g/L KH2PO4, 10 % v/v Hyclone Fetal Bovine Serum (ThermoFisher), heat-inactivated, 0.5% v/v Vitamin Mix (0.05 g D- biotin (Sigma), 0.05 g Folic acid (Sigma) and 2.5 g Sodium hydroxide) and Gentomycin (100 µg/mL), and incubated at 37°C in 75 cm^2^ tissue culture flasks (Iwaki, Japan). Total DNA was extracted from the cultures as previously described using the PowerSoil DNA Isolation kit (MO BIO Laboratories, USA) (Miller et al., 2015; Miller et al., 2017; Miller et al., 2018; Puzon et al., 2017). DNA was quantified using a Qubit (Invitrogen, USA) and stored at -80°C for subsequent genome sequencing.

*N. fowleri* LEE was grown axenically in oxoid media. Three replicates of *N. fowleri* LEE were passaged continuously through 50 B6C3F1 male mice. After mouse sacrifice, amoebae were extracted and grown in axenic media for one week to clear the culture of human cells prior to mRNA extraction.

*Genome sequencing and assembly*

*Naegleria fowleri* V212 DNA was prepared and sequenced as described in *Herman* et al., 2013 (PMID 23360210). Mitochondrial reads were first removed from 112,479,620 paired-end 100 bp Illumina and 609,044 454 reads using bowtie2 and the remaining reads were *de novo* assembled using SPAdes v3.1.1 (PMID 22506599) and scaffolded using SSPACE (PMID1149342), resulting in an assembly of 987 scaffolds >200bp totaling 27.7 Mb with an N50/N90of 92461/25477 bp and depth of coverage of 251X.

For *N. fowleri* 986, a genomic sequencing library was constructed using the Illumina Nextera sequencing library kit and sequenced on an Illumina Hiseq 2000 (100bp paired end sequencing). Raw sequence reads were imported into CLC Genomics (Qiagen), quality timed using the default parameters and assembled. Assembly statistics for V212, 986, and 30863 are provided inset below.

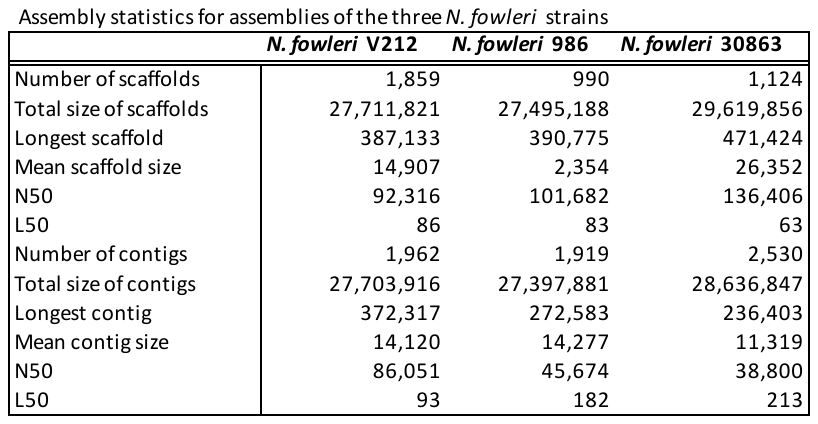

*Transcriptomics*

For gene prediction purposes, mRNA was extracted from *N. fowleri* CDC:V212 grown in axenic culture. Sequencing was done using an Illumina HiSeq platform, and ~150 million paired-end reads were generated. These were quality filtered using Trimmomatic (Bolger et al., 2014) and transcripts were *de novo* assembled using Trinity software (Haas et al., 2013) with default parameters. These transcripts were then used as hints to generate gene models using Augustus, described below.

For differential expression of genes associated with pathogenicity in *N. fowleri*, mRNA was extracted from three independent cultures of *N. fowleri* LEE-MP (mouse passaged), and from *N. fowleri* LEE-AX (grown only in axenic culture). One microgram RNA was converted to first strand cDNA in a 25 microliter volume (USB M-MLV RT). Twenty microliters (from 25 microliter total volume) from first strand synthesis was converted to double-strand cDNA (dscDNA; NEBNext mRNA Second Strand Synthesis Module, #E6111S).  Twenty microliters of dscDNA was then used for library generation (Illumina Nextera XT library preparation kit). Ampure beads were used for sample clean-up throughout. Libraries were sequenced on an Illumina MiSeq (2x300 cycles; 600V3 sequencing kit). Illumina MiSeq sequencing was performed at The Applied Genomics Centre at the University of Alberta, generating paired-end 2x300 reads. Reads were pre-processed using Trimmomatic v0.32400) (Bolger et al., 2014), by adaptor trimming, 5’ end trimming (15 bp), trimming of regions where the average Phred score is <20, and removal of short reads (<50 bp). Remaining read set quality and characteristics were visualized using FastQC v0.11.2.401 (Andrews, 2010).

The *N. fowleri* LEE-AX and LEE-MP reads were mapped to the *N. fowleri* V212 genome using TopHat v2.0.10 (Kim et al., 2013), with minimum intron length set to 30bp, based on an assessment of predicted genes. Transcripts were then generated using Cufflinks with the Reference Annotation Based Transcript option, using the predicted genes as a reference dataset (Roberts et al., 2011; Trapnell et al., 2010). The reference transcripts are tiled with ‘faux-reads’ to aid in the assembly, and these sequences are added to the final dataset containing the newly assembled transcripts. In order to obtain transcripts not represented by genes in the *N. fowleri* V212 genome, Trinity (release 2013-02-25) was used for purely *de novo* transcriptome assembly, using a genome-guided approach, with a --genome_guided_max_intron value of 5000(Haas et al., 2013; Trapnell et al., 2012). Novel Trinity-generated transcripts were added to the Cufflinks-generated transcriptome prior to downstream differential expression analyses.

*Differential expression analysis*

Differential expression (DE) analyses were performed for the pathogenicity transcriptomic data using the programs Cuffdiff (Trapnell et al., 2012) and Trinity (Haas et al., 2013). Cuffdiff was first used to map reads to the final transcriptome assembly, with high-abundance (>10,000 reads) mitochondrial and extrachromosomal plasmid genes masked, and differential expression was calculated using geometric normalization. The Trinity Perl-to-R (PtR) toolkit was used to assess variation between replicates. One mouse passage replicate, MP2, was highly dissimilar to other mouse passaged and axenically grown samples, and was therefore excluded from further analysis. Using the Trinity suite of scripts, reads were aligned and transcript abundance estimated using RSEM (Li and Dewey, 2011) with TMM library normalization. Differentially expressed transcripts were then identified using EdgeR (Robinson et al., 2010). Comparison of these transcripts with those identified by Cuffdiff revealed highly overlapping datasets, with only a few genes considered to be differentially expressed by EdgeR but not Cuffdiff. Because this work is exploratory and we sought to minimize the potential for false negatives, we chose a lax false discovery rate cutoff of 0.1. However, the majority of up-regulated genes have FDR values less than 0.05.

*Gene prediction and annotation*

Gene prediction was performed using the program Augustus v.2.5.5 (Stanke and Waack, 2003; Stanke et al., 2004), incorporating the HiSeq-generated transcript dataset as extrinsic evidence termed “hints”. Furthermore, Augustus was also trained using a manually annotated 60kb segment from the *N. fowleri* V212 genome published previously (Herman et al., 2013). This region was annotated by using it as a TBLASTN query to search the non-redundant database. Gene boundaries were identified using the alignments of the top hits, and genes were annotated based on top BLAST hit identities. Gene prediction was performed for the *N. fowleri* CDC:V212 and 986 genomes generated by this study, as well as on the publicly available genome for strain ATCC ATCC30863. For each genome, the parameter --alternatives-from-evidence was set to true, as this reports alternative gene transcripts if there is evidence for them (i.e. from transcriptome dataset). The parameter --alternatives-from-sampling was also set to true, as this outputs additional suboptimal transcripts. Parameters for determining the importance of different hint data were kept as default.
Annotation of genes in specific subsystems of focus in this manuscript was done using homology searching. Functionally characterized homologues of proteins of interest were used as BLASTP (Altschul et al., 1990) queries to search the predicted proteins from the *N. fowleri* strain genomes. At a minimum, putative *N. fowleri* homologues must be retrieved with an E-value <0.05. To be considered true homologues, the *N. fowleri* protein must retrieve the original query or a clear orthologue in a reciprocal BLAST search also with an E-value <0.05.

*Phylogenetic analysis*

Bayesian and Maximum-Likelihood phylogenetics analyses were performed to assign orthology to proteins from highly paralogous families. Sequences were aligned using MUSCLE v.3.8.31(Edgar, 2004), alignments were visualized in Mesquite v.3.2 (Maddison and Maddison, 2015), and manually masked and trimmed to remove positions of uncertain homology. ProtTest v3.4(Darriba et al., 2011) was used to determine the best-fit model of sequence evolution. Phylobayes v4.1 (Lartillot et al., 2009) and MrBAYES v3.2.2 (Ronquist et al., 2012) programs were run for Bayesian analysis and RAxML v8.1.3 (Stamatakis, 2014) was run for Maximum-Likelihood analysis. Phylobayes was run until the largest discrepancy observed across all bipartitions was less than 0.1 and at least 100 sampling points were achieved, MrBAYES was used to search treespace for a minimum of one million MCMC generations, sampling every 1000 generations, until the average standard deviation of the split frequencies of two independent runs (with two chains each) was less than 0.01. Consensus trees were generated using a burn-in value of 25%, well above the likelihood plateau in each case. RAxML was run with 100 pseudoreplicates.

*BUSCO analysis*

BUSCO v3 software (Simão et al., 2015; Waterhouse et al., 2018) was used to assess genome completeness. Predicted proteins were used as input, with the eukaryote dataset eukaryota_odb9 set of Hidden Markov Models.

*Synteny analysis*

The program Mauve (build 2015-02-25) (Darling et al., 2004; Darling et al., 2010) was used with the progressiveMauve alignment method to visualize synteny across the three *N. fowleri* genomes and the *N. gruberi* genome. Because the genomes being aligned are from relatively closely related taxa, the match seed weight of 15 was used. Both ‘Full Alignment’ and ‘Iterative Refinement’ parameters were selected. These parameters direct Mauve to first perform a recursive anchor search and full genome alignment using MUSCLE, followed by guide tree-independent refinement of the alignment. Default gap open and gap extend scores of -400 and -30 were used, respectively, as was the suggested HOXD scoring matrix.

*Orthologous groups analysis*

To identify orthologous groups of sequences between the four *Naegleria* predicted proteomes, the program OrthoMCL v.2.0.9424 was used (Li et al., 2003). The Markov Clustering algorithm is an unsupervised clustering algorithm that clusters graphs based on pairwise scores (in this case, normalized E-values following an all-versus-all BLAST) and an inflation value. This latter value controls the clustering tightness, and for these analyses, was kept at the suggested 1.5.

*Trans-membrane domain prediction*

The program TMHMM v2.0426 (Krogh et al., 2001; Sonnhammer et al., 1998) was used to detect trans-membrane helices in all *N. fowleri* V212 proteins. This software uses an HMM generated from 160 cross-validated membrane proteins, and outputs all predicted helices and protein orientation in the membrane. Because trans-membrane helix prediction was done to generate a list of potential G-protein coupled receptors investigated further by domain prediction, scoring cutoffs were not used.

*Cytoskeletal protein annotations*

To assess the cytoskeletal repertoire of the three *Naegleria fowleri* isolates, we first compared *N. fowleri* proteins to previously identified *N. gruberi* actin and microtubule associated proteins (Fritz-Laylin et al., 2010) using BLAST (Altschul et al., 1990) against the protein databases generated in Augustus for each of the three *N. fowleri* isolates, using default search parameters. The top *N. fowleri* hits identified by comparisons to *N. gruberi* were then compared using BLAST to the full *N. gruberi* protein library to establish best mutual BLAST hits, which are included in Supplementary Table 5. We then searched for additional proteins not found in *N. gruberi* using human or *Dictyostelium discoidium* protein sequences obtained from pubmed and dictyBase (http://dictybase.org), respectively*.* To identify *N. fowleri* homologs, we again used BLAST with the default parameters except replacing the scoring matrix with the BLOSUM45 scoring matrix. We further validated protein identities through hmmscan searches using a gathering threshold and Pfam domains using the HMMER website (https://www.ebi.ac.uk/Tools/
hmmer/search/hmmscan).
